# Network Adaptability Governs Resilience and Susceptibility to Social Defeat

**DOI:** 10.1101/2025.11.26.690729

**Authors:** Sarah Ayash, Stefan Paul Koch, Jeehye An, Ulrich Schmitt, Susanne Mueller, Marco Foddis, Esin Candemir, Oliver Tuescher, Philipp Boehm-Sturm, Marianne Mueller

## Abstract

Stress resilience is defined as the ability to maintain mental health after adversity. In humans, resilience is marked by threat–safety discrimination and extinction of fear, whereas susceptibility involves fear generalization and extinction resistance. Using a translational mouse model of chronic social defeat stress combined with preregistered multimodal brain imaging, we investigated structural and network-level markers of resilience, susceptibility, and threat learning. Resilience was characterized by preserved dentate gyrus integrity, greater microstructural complexity in prefrontal regions, and signatures consistent with enhanced inhibitory gating within amygdalar circuits. Susceptibility, in contrast, involved reduced dentate gyrus complexity and diminished microstructural flexibility, with a weaker but parallel pattern in the basolateral amygdala. Impaired threat learning was linked to compromised CA1/CA2 integrity and reduced pons connectivity, highlighting hippocampal–brainstem interactions in memory consolidation. Overall, these findings show that resilience emerges from adaptive network reorganization, whereas susceptibility and impaired learning reflect distinct dysfunctions, underscoring individual coping differences even among genetically identical animals.

## INTRODUCTION

Resilience is defined as the maintenance or rapid recovery of mental health when facing adversity (1). In humans, two key behavioral features are associated with resilience: the ability to discriminate between threat and safety cues, and the ability to extinguish fear when danger has passed. In contrast, susceptibility is marked by fear generalization and resistance to extinction (2). Rodent models, particularly chronic social defeat (CSD) in male mice, have been instrumental in advancing our understanding of resilience (3). Some mice develop social avoidance after repeated defeats, and we showed that this avoidance reflects correct conditioned learning where the aggressors’ strain is the threat-associated cue (4). Using our validated social threat-safety test (STST), we recently identified three distinct subgroups after CSD: discriminating-avoiders (avoid only the aggressors’ strain, i.e., threat-safety discrimination), indiscriminate-avoiders (generalize fear to safe strains), and non-avoiders (fail to learn the threat association; 5–6; Figure 1A). Only discriminating-avoiders extinguish avoidance (when facing aggressors’ strain from a safe distance), consistent with resilience in humans, whereas indiscriminate-avoiders resist extinction, reflecting susceptibility. Non-avoiders represent a third group with impaired aversive learning. The ability to distinguish aversive from safe cues is critical for survival, and because this learning depends on the fear circuitry—one of the most conserved neural systems across species—our model provides a powerful framework for translational investigation. We previously showed distinct transcriptional signatures across the three subgroups in basolateral amygdala (BLA), medial prefrontal cortex (mPFC), and ventral hippocampus (vHC; 5). Other brain regions implicated in defeat-induced avoidance include the dorsal HC (dHC; 7), and the ventrolateral subdivision of the ventromedial hypothalamus, part of the hypothalamic medial zone (HMZ; 8). The fact that robust differences in behavior and underlying biology still emerge in mice despite negligible genetic variability and highly uniform environmental conditions underscores the fundamental individuality of stress coping.

**Figure 1.**
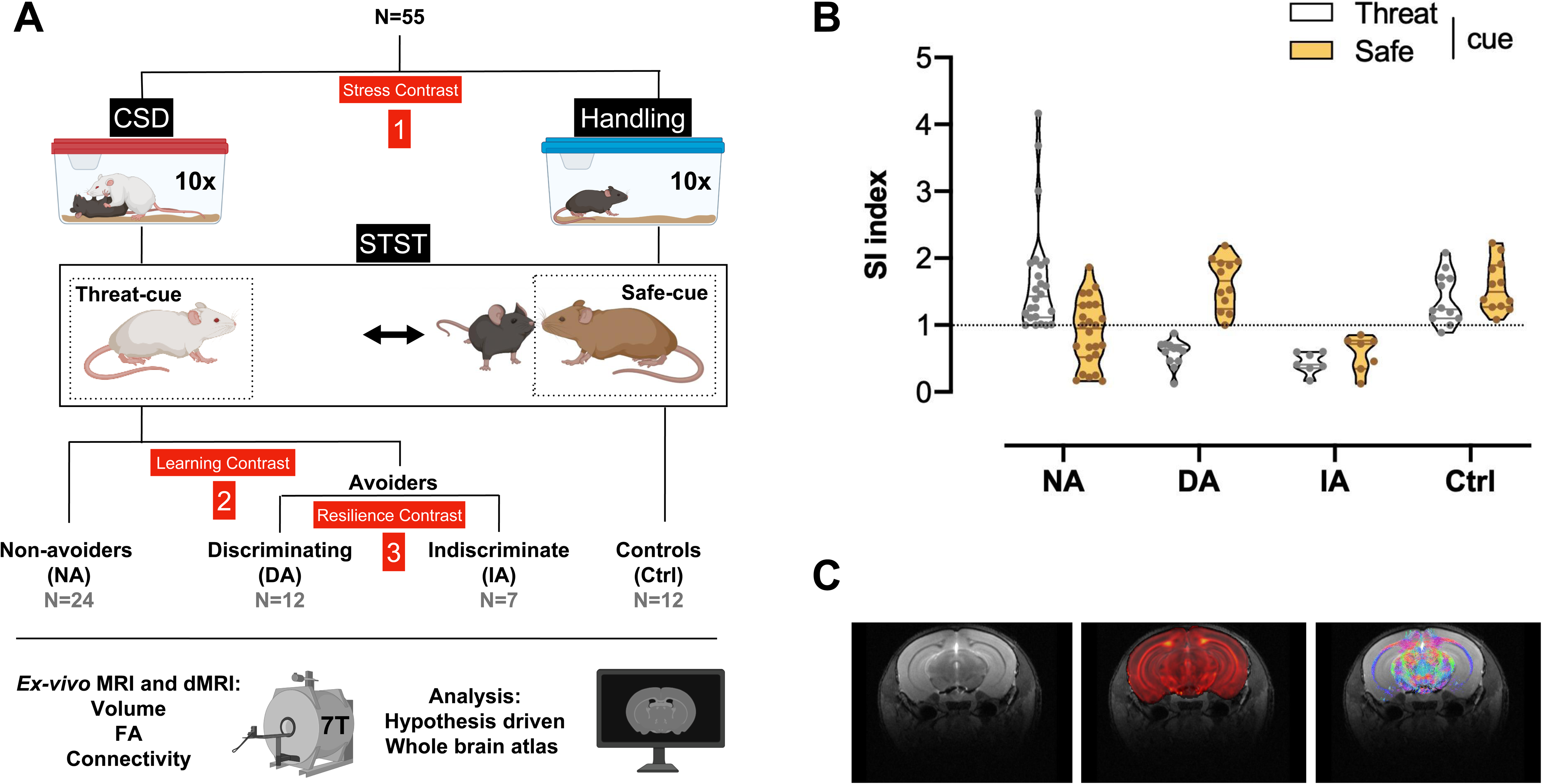
Three defeated subgroups. (A) Animal model and conceptual figure. Chronic social defeat (CSD) or handling took place for 10 days and were followed by social threat-safety test (STST) a day later, which in turn was immediately followed by perfusion. During STST, two conspecifics are presented, one belongs to the threat-associated strain (cue) from the CSD days, the other to the unknown 129/Sv safe strain. Unlike avoiders, non-avoiders (NA) don’t avoid the threat cue. Within the avoiders, indiscriminate-avoiders (IA) also avoid the safe cue whereas discriminating-avoiders (DA) don’t. Numbers in red boxes represent the three analyses allowing assessment of differences associated with 1. Stress: defeated versus handled controls. 2. Threat learning: NA versus avoiders of threat. 3. Resilience: DA versus IA. Animals were brain imaged using 7 Tesla scanner with cryoprobe. Region-wise volume, diffusion magnetic resonance imaging (dMRI) fractional anisotropy (FA), and dMRI connectivity were obtained. Hypothesis-driven analysis on 10 brain regions was performed to identify differences in volume and FA while exploratory analysis on whole brain atlas level was performed to dissect connectivity differences. (B) STST. NA display a social interaction index ≥1 with the threat cue (NA; n=24), DA display a social interaction index ≥1 exclusively with the safe cue (DA; n=12), and IA display a social interaction index <1 with both strains (IA; n=7). Non-defeated controls (Ctrl; n=12) have comparable indices with both strains. Results are presented as truncated violin plots. Each animal is represented by two data points, one with each cue. (C) Representative brain image. Representative examples of T2-weighted anatomical MR images to assess volumetric changes (left), FA maps reconstructed from dMRI as a marker of microstructural changes (middle, FA in red overlaid on grayscale anatomical image) and dMRI whole brain connectome to measure changes in structural connectivity between brain regions (right, reduced set of fibers with color-encoded directionality overlaid on anatomical image).

While transcriptional studies revealed subgroup-specific molecular differences, no previous work has examined structural or network-level alterations using brain imaging in these behaviorally defined subgroups. This represents a critical next step: imaging is not only feasible in rodents but directly translatable to humans, enabling the identification of imaging-based markers of resilience, susceptibility, and impaired aversive learning. Importantly, brain imaging offers a brain-wide, integrative view of neural organization that neither molecular nor circuit-level approaches alone can provide. By capturing coordinated changes across distributed systems in a hypothesis-free manner, imaging allows us to uncover large-scale structural and connectivity patterns that would otherwise remain undetected. To investigate structural and connectivity changes underlying these behavioral phenotypes, we applied magnetic resonance imaging (MRI) and diffusion MRI (dMRI) to CSD-exposed mice phenotyped with the STST. We analyzed three contrasts: 1) stress exposure (defeated vs. control), 2) threat learning (non-avoiders vs. avoiders), and 3) resilience (discriminating-vs. indiscriminate-avoiders; Figure 1A). By additionally comparing each subgroup after defeat to the non-defeated control group, we were able—albeit with some limitations—to distinguish which differences between subgroups reflect mal/adaptations to chronic stress exposure and which represent features maintained during the stress experience. We quantified volume differences across 10 brain regions of interest (ROI) previously implicated in stress responsiveness—BLA, vHC, CA1/2, dentate gyrus (DG), infralimbic (IL), prelimbic (PL), anterior cingulate cortex (ACC), HMZ, nucleus accumbens (NAc), and ventral tegmental area (VTA). We then used dMRI to assess fractional anisotropy (FA), explore whole-brain connectivity, and apply graph theory analyses to quantify network topology, namely clustering coefficient (local segregation) and local efficiency (local integration; 9).

Importantly, **we preregistered this study** in the Open Science Framework (https://doi.org/10.17605/OSF.IO/MDTH4) **before conducting any brain imaging analysis**—a critical step to enhance transparency, prevent data dredging, and strengthen the robustness and reproducibility of our findings. These issues are particularly acute in brain imaging, where analytic flexibility, high-dimensional data, and variable preprocessing pipelines make studies especially vulnerable to false positives and poor reproducibility. Preregistration therefore represents a critical safeguard against these issues. To our knowledge, this is the first preregistered study combining brain imaging with CSD paradigm.

Our findings reveal that resilience involves preserved DG integrity, greater microstructural complexity in prefrontal regions, unique connectivity patterns—particularly in the retrohippocampal region (RHR) and features consistent with enhanced inhibitory gating in amygdalar circuits. By contrast, susceptibility is marked by diminished DG complexity and reduced microstructural flexibility, with a similar but less pronounced pattern in the BLA. Impaired aversive learning aligns with compromised CA1/CA2 integrity and reduced pons connectivity, highlighting the contribution of hippocampal–brainstem interactions to threat-memory consolidation. Additionally, chronic stress exposure itself is associated with reduced prefrontal volume and distinct connectivity alterations, most prominently weaker coupling in the pallidum, underscoring the broader network-level impact of prolonged social stress.

Several studies have applied MRI or dMRI in the CSD model (10–18). However, none have examined brain structure and network-level markers in relation to *translationally* defined behavioral features of resilience (threat-safety discrimination and responsiveness to extinction). Here, we aimed to combine high-resolution MRI, dMRI, and graph theory analyses with a model that is both translationally relevant to humans and ecologically valid in animals (2–3). This systems-level approach represents the first application of brain imaging to these specific phenotypes and provides a framework for identifying imaging-based markers of individual differences in stress coping.

## RESULTS

### Stratification of defeated animals based on threat learning and threat-safety discrimination

STST conducted after CSD identifies three subgroups based on the calculation of two social interaction (SI) indices, one with each strain/cue (Figure 1A-B). Non-avoiders interact with the mice from the threat-associated strain (SI index ≥1). In the group of the avoiders of the threatening strain, two behavioral phenotypes exist; discriminating-avoiders that maintain social interaction exclusively with the safe strain (SI index ≥1; thereafter referred to as resilient group) and indiscriminating-avoiders that also avoid the safe strain (SI index <1). Such fear generalization is the hallmark of the susceptible group. Non-defeated controls show comparable indices with both strains.

### Smaller volumes in stressed and susceptible animals

We next measured the volume of several brain structures implicated in the stress network. We found smaller volumes in mice that underwent CSD compared to non-stressed controls in subregions of the mPFC, namely the ACC and PL (Figure 2A). Meanwhile, the susceptible phenotype (indiscriminate-avoiders) was associated with smaller volumes compared to resilience (discriminating-avoiders) in the vHC, DG, and NAc (Figure 2B and Supplementary Fig. 1). Compared to non-stressed controls, indiscriminate-avoiders displayed smaller volumes in DG and NAc while discriminating-avoiders displayed comparable volumes (Figure 2B, Supplementary Fig. 1, and Supplementary Table 1). Interestingly, threat learning (as assessed by comparing non-avoiders of threat to the avoiders group) was not associated with any volumetric differences.

**Figure 2.**
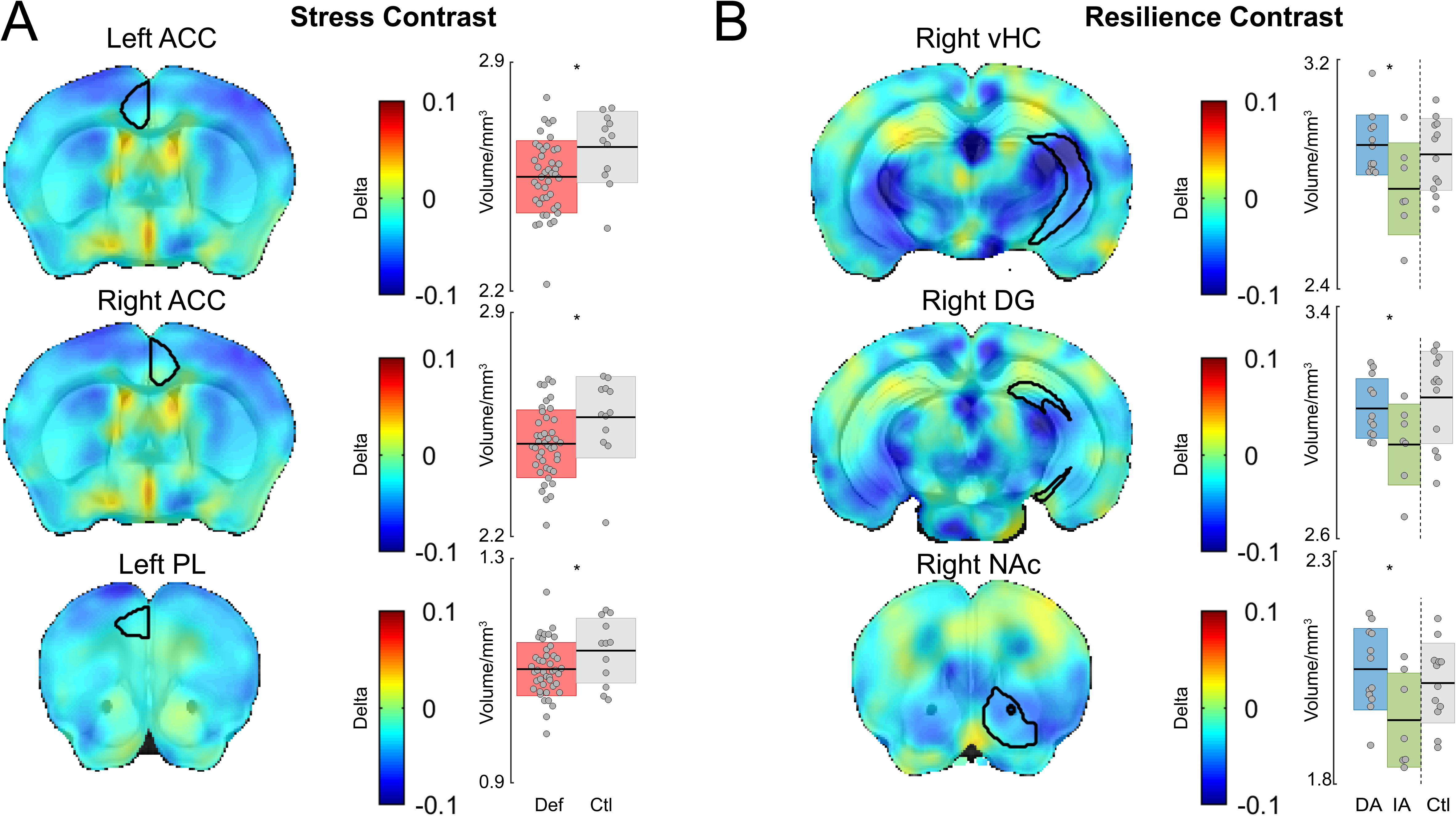
Region wise volume. (A) Chronic stress exposure-associated differences. Subregions of the medial prefrontal cortex (mPFC), namely the anterior cingulate cortex (ACC), and the pre-limbic cortex (PL) are significantly smaller in the defeated group of animals (Def; n=43) compared to the controls (Ctl; n=12). (B) Resilience-associated differences. Ventral hippocampus (vHC) and nucleus accumebns (NAc) are significantly larger in resilient discriminating-avoider (DA; n=12) compared to susceptible indiscriminate-avoider (IA; n=7). Data from control animals is displayed for visualization purposes only and was excluded from statistical analysis. Brain maps depict the differences quantified in the respective box plot, the region of interest is marked in black, color bars indicate relative volume differences, with volumes normalized to the Allen Brain Atlas and compared between groups. Box plots show mean±SD., t-test, two-tailed, ACC left hemisphere (p-value=0.01, Mean of Def=2.55 and SD=0.11, Mean of Ctl=2.64 and SD=0.11, t-value=-2.53, df=53), ACC right hemisphere (p-value=0.03, Mean of Def=2.49 and SD=0.11, Mean of Ctl=2.57 and SD=0.13, t-value=-2.29, df=53), PL (p-value=0.05, Mean of Def=1.10 and SD=0.05, Mean of Ctl=1.14 and SD=0.06, t-value=-2.03, df=53), vHC (p-value=0.02, Mean of DA=2.90 and SD=0.10, Mean of IA=2.75 and SD=0.16, t-value=2.53, df=17), DG (p-value=0.04, Mean of DA=3.05 and SD=0.10, Mean of IA=2.92 and SD=0.14, t-value=2.23, df=17), and NAc (p-value=0.02, Mean of DA=2.05 and SD=0.09, Mean of IA=1.94 and SD=0.10, t-value=2.45, df=17).

### Larger fractional anisotropy in susceptible animals

Although FA is typically higher in white matter, measurable anisotropy in gray matter regions provides meaningful indices of local microstructural organization. Threat learning was associated with a difference in FA exclusively in the CA1/2 region of the HC. Smaller FA was observed in the group failing to learn to avoid threat (non-avoiders) compared to avoiders of threat (Figure 3A). Compared to the non-stressed control group, neither of the non-avoiders or avoiders groups were different (Figure 3A). The largest number of differences in FA (6 out of the 10 ROI) was between resilient and susceptible animals (discriminating-vs. indiscriminate-avoiders, respectively). Discriminating-avoiders were associated with smaller FA compared to indiscriminate-avoiders. Here, DG, BLA, HMZ, and subregions of the mPFC, namely the ACC, PL, and IL, all displayed smaller FA in discriminating-avoiders (Figure 3B). Compared to non-stressed controls, indiscriminate-avoiders displayed larger FA in DG and BLA (Figure 3B and Supplementary Table 2A). By contrast, compared to non-stressed controls discriminating-avoiders displayed smaller FA in PL and IL (Figure 3B and Supplementary Table 2B). Neither of the two groups were significantly different to the controls in HMZ and ACC. Of note, chronic stress exposure was not associated with any differences in FA.

**Figure 3.**
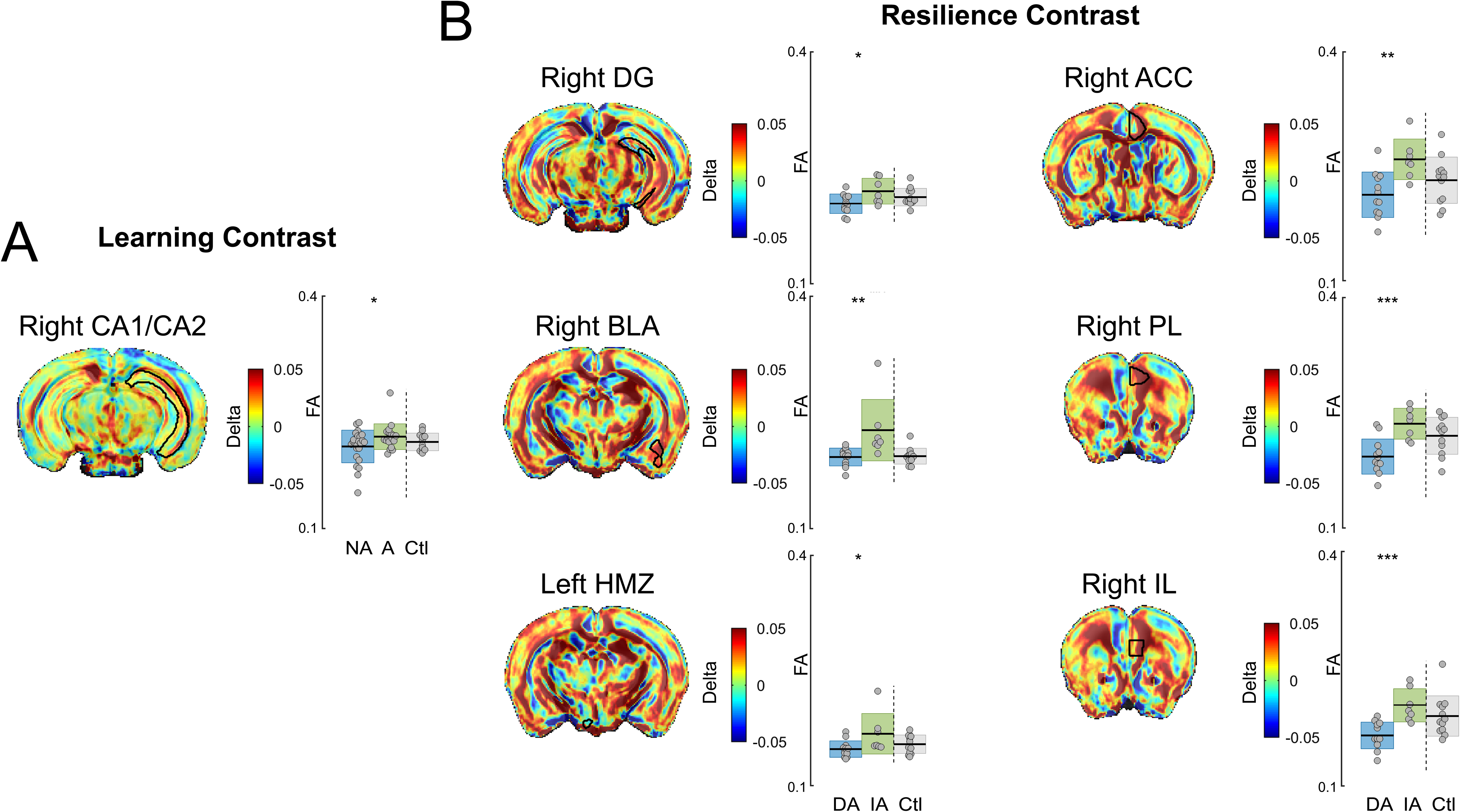
Diffusion magnetic resonance imaging fractional Anisotropy (FA). (A) Threat learning-associated differences. Animals that don’t conditionally learn to avoid the threatening strain (non-avoider; NA; n=24) display significantly lower FA in the CA1/CA2 of the hippocampus compared to avoiders of threat (A; n=19). (B) Resilience-associated differences. Dentate gyrus (DG), basolateral amygdala (BLA), hypothalamic medial zone (HMZ), anterior cingulate cortex (ACC), pre-limbic area (PL), and infra-limbic area (IL) all have significantly lower FA in resilient discriminating-avoiders (DA; n=12) compared to susceptible indiscriminate-avoiders (IA; n=7). Data from control animals is displayed for visualization purposes only and was excluded from statistical analysis. Brain maps depict the differences quantified in the respective box plot, the region of interest is marked in black, color bars indicate relative FA differences, with values normalized to the Allen Brain Atlas and compared between groups. Box plots show mean±SD, t-test, two-tailed, (A) CA1/2 (p-value=0.04, Mean of NA=0.21 and SD=0.02, Mean of avoiders=0.22 and SD=0.02, t-value=-2.14, df=41), (B) DG (p-value=0.03, Mean of DA=0.20 and SD=0.01, Mean of IA=0.22 and SD=0.02, t-value=-2.39, df=17), BLA (p-value=0.01, Mean of DA=0.19 and SD=0.01, Mean of IA=0.23 and SD=0.04, t-value=-2.88, df=17), HMZ (p-value=0.03, Mean of DA=0.15 and SD=0.01, Mean of IA=0.17 and SD=0.03, t-value=-2.34, df=17), ACC (p-value<0.01, Mean of DA=0.21 and SD=0.03, Mean of IA=0.26 and SD=0.03, t-value=-3.40, df=17), PL (p-value<0.01, Mean of DA=0.19 and SD=0.02, Mean of IA=0.24 and SD=0.02, t-value=-4.10, df=17), and IL (p-value<0.01, Mean of DA=0.16 and SD=0.02, Mean of IA=0.20 and SD=0.02, t-value=-4.35, df=17).

### Largest number of differential connectivity patterns is in resilient animals

We also investigated dMRI connectivity, where we used an unbiased approach (whole brain atlas of 44 regions) to ascertain connectivity differences of circuits and regions not previously implicated in stress, threat learning, and resilience. By employing dMRI, we instructed probable diffusion tracts to obtain information on brain connectivity. Only connections that are within the same hemisphere and appear by viral tracing (based on the Allen mouse brain connectivity atlas) were considered.

Chronic stress exposure was associated with the smallest number of different patterns in connectivity (24 patterns) compared to threat learning (37 patterns) and resilience-associated differences (56 patterns; Figure 4 and Supplementary Table 3). Resilience-associated differences (as identified by comparing discriminating-avoiders to indiscriminate-avoiders) involved the largest number of regions (34 regions), followed by threat learning (non-avoiders vs. avoiders; 29 regions), and lastly, chronic stress exposure (defeated vs. controls; 23 region; Figure 4 and Supplementary Table 3). Comparable number of stronger and weaker differential connectivity patterns were identified in every sub/group comparison (Figure 4). The largest number of different connectivity patterns between defeated and non-stressed controls was in the pallidum (weaker connectivity in defeated), between non-avoiders of threat and avoiders in pons (weaker in non-avoiders), and between resilient discriminating-avoiders and susceptible indiscriminate-avoiders in the RHR (large majority weaker in discriminating-avoiders; Figure 4 and Supplementary Table 3).

**Figure 4.**
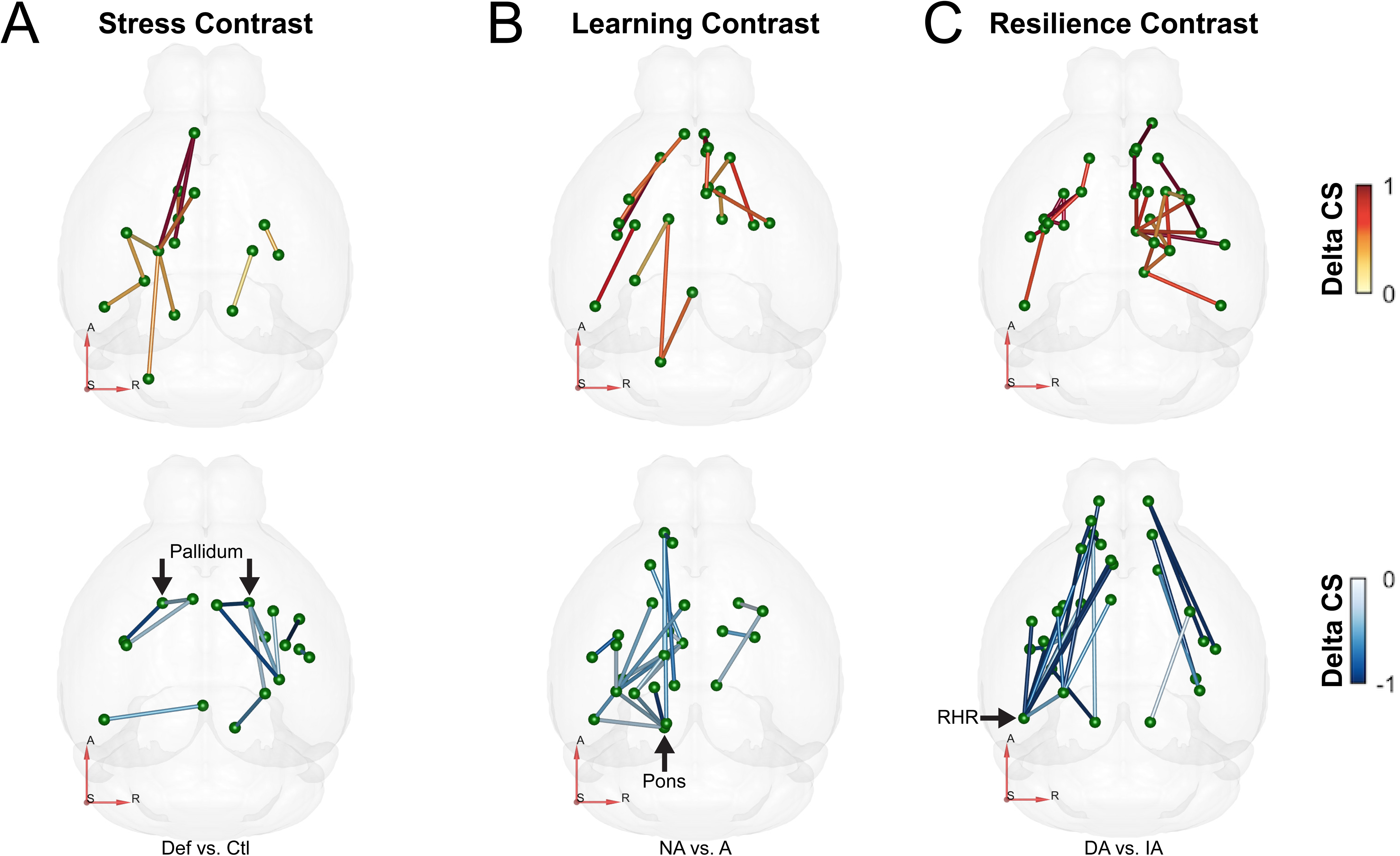
Diffusion magnetic resonance imaging connectivity. (A) Chronic stress exposure-associated differences. A total of 24 differential connectivity patterns involving 29 brain regions (irrespective of hemispheres) are identified between defeated animals (n=43) and handled controls (n=12). Out of the 24, 13 are less connected in defeated animals (lower panel) (B) Threat learning-associated differences. A total of 37 differential connectivity patterns involving 39 brain regions (irrespective of hemispheres) are identified between animals that don’t conditionally learn to avoid the threatening strain (non-avoider; n=24) and avoiders (n=19) of threat. Out of 37, 22 are less connected in non-avoiders (lower panel). (C) Resilience-associated differences. A total of 56 differential connectivity patterns involving 47 brain regions (irrespective of hemispheres) are identified between resilient discriminating-avoiders (n=12) and susceptible indiscriminate-avoiders (n=7). Out of the 56, 28 are less connected in discriminating-avoiders (lower panel). Group differences in connectivity strength (Delta CS, unitless), RAS (right, anterior, superior) coordinate system axes shown in the left lower corner of each subfigure in red. Highlighted brain regions in each panel represent those showing the greatest number of connectivity-pattern changes, irrespective of whether these alterations were predominantly up- or down-regulated. RHR stands for retrohippocampal region.

### Larger clustering coefficient and effeciency in the DG of susceptible and IAN of resilient animals

To integrate our findings with neurobiologically meaningful and easily computable measures in a brain-wide neural connectivity, we used unbiased graph theory analysis (9). We represented the brain as network (graph) of 44 brain regions (nodes) and connections (edges). We investigated **clustering coefficient and local network efficiency**. Clustering coefficient is a measure of how well regions connected to a node are also highly connected among each other. It is a measurement of *local segregation* in the brain, which is the ability for specialized processing to occur within densely interconnected groups of brain regions (9). Meanwhile, local efficiency reflects how efficiently information is exchanged within those local clusters. It is a measurement of *local integration*, which is the ability to rapidly combine specialized information from distributed brain regions (9).

Significant findings were associated with chronic stress exposure and resilient phenotype but not with aversive conditioned learning. Specifically, accessory olfactory bulb (AOB) displayed a lower clustering coefficient and local efficiency in defeated animals compared to controls (Figure 5A). Meanwhile, clustering coefficient of substantia nigra compact part (SN) and IAN as well as local efficiency were higher in resilient discriminating-avoiders compared to susceptible indiscriminate-avoiders (Figure 5B). Compared to non-defeated controls, resilient discriminating-avoiders’ values were significantly greater in IAN (Figure 5B and Supplementary Table 4A-B). In contrast, DG showed the opposite i.e. lower clustering coefficient and local efficiency in the resilient group compared to the susceptible one. Compared to non-defeated controls, indiscriminate-avoiders displayed significantly larger local efficiency in DG, a similar trend was observed in clustering coefficient (Figure 5B and Supplementary Table 4C-D).

**Figure 5.**
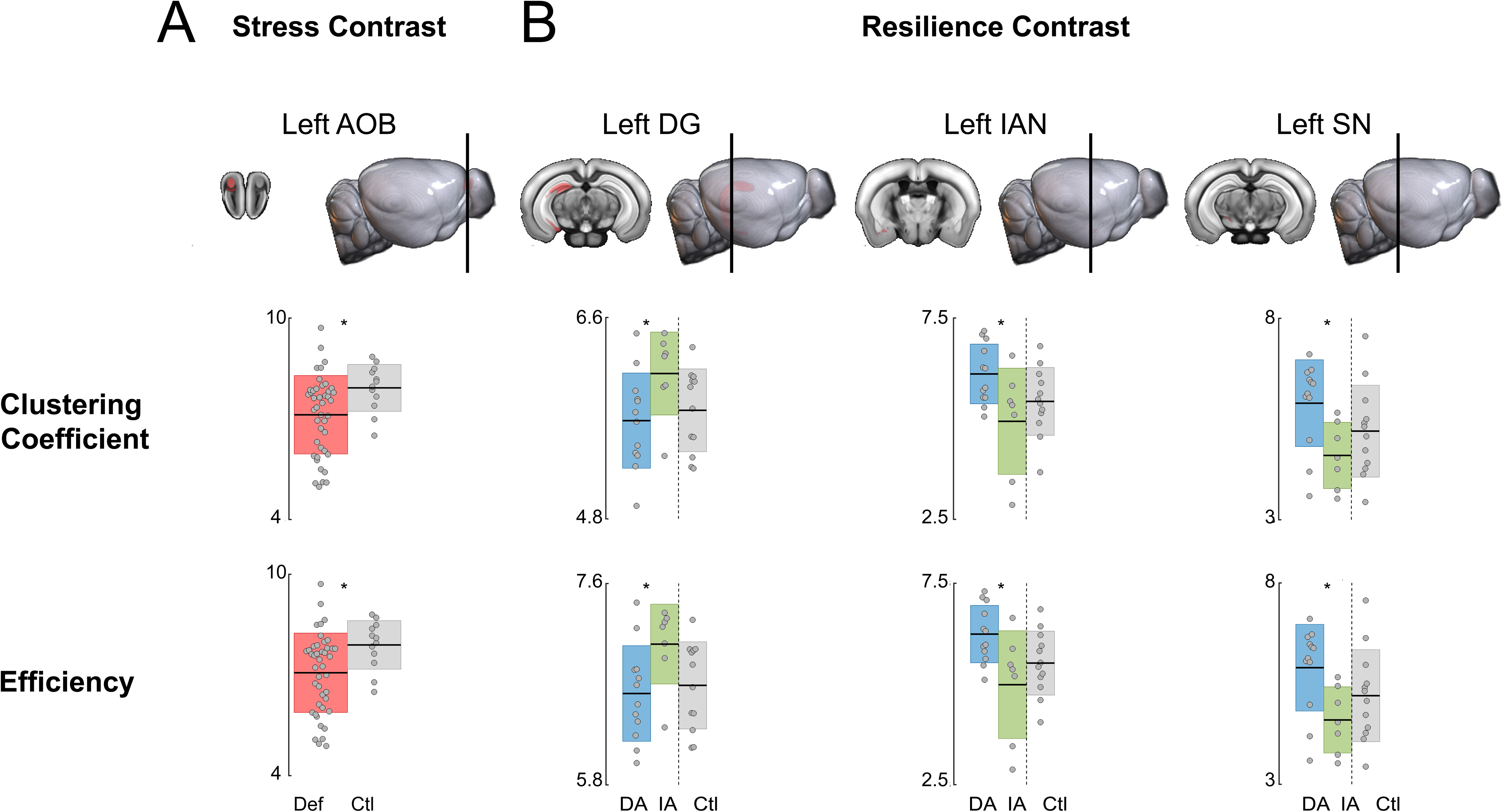
Graph theory: clustering coefficient and local efficiency. (A) Chronic stress exposure-associated differences. Accessory olfactory bulb (AOB) displays a lower clustering coefficient and local efficiency in defeated animals (Def; n=43) compared to controls (Ctl; n=12). (B) Resilience-associated differences. Substantia nigra compact part (SN) and intercalated amygdala nucleus (IAN) clustering coefficient and local efficiency are higher in resilient discriminating-avoiders (DA; n=12) compared to susceptible indiscriminate-avoiders (IA; n=7). In contrast, DG shows lower clustering coefficient and local efficiency in IA. Data from control animals is displayed for visualization purposes only and was excluded from statistical analysis. mean±SD, t-test, two-tailed, (A) AOB clustering coefficient (p-value=0.03, Mean of Def=7.06 and SD=1.18, Mean of Ctl=7.89 and SD=0.72, t-value=-2.30, df=53) and local efficiency (p-value=0.03, Mean of Def=7.12 and SD=1.17, Mean of Ctl=7.92 and SD=0.70, t-value=-2.26, df=53). (B) SN clustering coefficient (p-value=0.01, Mean of DA=5.89 and SD=1.08, Mean of IA=4.60 and SD=0.82, t-value=2.75, df=17), IAN (p-value=0.02, Mean of DA=6.11 and SD=0.74, Mean of IA=4.93 and SD=1.32, t-value=2.51, df=17), as well as local efficiency (SN: p-value=0.01, Mean=5.89 and SD=1.08, Mean=4.60 and SD=0.82, t-value=2.75, df=17; IAN: p-value=0.02, Mean=6.24 and SD=0.71, Mean=4.98 and SD=1.34, t-value=2.70, df=17), and DG clustering coefficient (p-value=0.04, Mean=5.68 and SD=0.43, Mean=6.10 and SD=0.37, t-value=-2.18, df=17) and local efficiency (p-value=0.03, Mean=6.62 and SD=0.42, Mean=7.06 and SD=0.36, t-value=-2.30, df=17).

## DISCUSSION

In this preregistered study, we set out to determine whether structural and network-level brain markers can be identified for resilience, susceptibility, and impaired aversive learning. Using a translationally valid mouse model of CSD with our validated STST combined with multimodal brain imaging, we specifically asked 1. how chronic social stress alters brain structure and connectivity, 2. how the underlying biology of impaired threat learning differs from successful conditioned avoidance, and 3. how resilience and susceptibility can be distinguished at the neurobiological level. We found distinct volumetric, FA, connectivity, and topological differences between the different phenotypes.

A central finding on how **chronic social stress exposure** alters brain structure involves the mPFC, a region critical for regulating hypothalamic-pituitary-adrenal axis activity under stress (19–20) and consistently reported as reduced in PTSD patients (21–23). We found smaller mPFC subregions (ACC and PL) in defeated animals compared to controls. Such reductions may result from dendritic retraction and spine loss (24). Connectivity analyses revealed that the pallidum displayed the largest number of different connectivity patterns between defeated and control animals. Specifically, weakened connections were found in the defeated. The pallidum is a multifunctional limbic hub involved in motivation, reward processing, and arousal regulation, and also contributes to sleep–wake control (25–26). It maintains strong connectivity with frontal cortical regions in both rodents (27) and humans (28). Thus, altered pallidum–ACC coupling in defeated animals may reflect stress-related disturbances in circuits that integrate motivational, affective, and arousal signals (29). These findings are consistent with prior imaging work showing weaker ACC connectivity (16) and altered PFC tissue properties following chronic social defeat (17).

**Impaired aversive learning** in non-avoiders was associated with lower FA in CA1/CA2 and weakened connections in the pons. The locus coeruleus, a noradrenergic hub within the pons, modulates hippocampal activity during learning, and reduced pons–hippocampal connectivity may therefore impair communication within this memory network (30). Given the critical role of both the hippocampus and pons in memory consolidation, these findings suggest that the failure of this group to acquire threat-associated avoidance reflects disrupted consolidation processes.

The HC consistently emerged as a central hub differentiating resilient from susceptible animals. **Susceptible mice** displayed smaller DG volume relative to both resilient and control groups. Stress is known to reduce hippocampal volume (10, 31), while antidepressants promote neurogenesis (32). This pattern extended to microstructural measures. Susceptible animals showed higher FA across 6 out of the 10 ROI, including the DG. Because FA values in gray matter are normally low due to the complex and multidirectional arrangement of cell bodies, dendrites, and small-caliber axons, elevated FA in susceptible animals likely reflects microstructural simplification—such as dendritic retraction, loss of cellular diversity, or reduced orientation variability—resulting in fewer diffusion barriers and diminished microstructural complexity. At the network level, susceptibility was marked by increased network clustering and efficiency in the DG. Connectivity analyses reinforced this picture. The RHR, which provides contextual and temporal input to the HC relevant for threat–safety discrimination (34), showed the largest number of different connectivity patterns between resilient and susceptible mice, with a majority of stronger connections in the latter. This DG profile—smaller volume, higher FA, increased network clustering/efficiency—paints a consistent picture of stress-induced atrophy and a less adaptable DG network, potentially reducing its capacity for experience-dependent plasticity and aligning with impaired discrimination and fear generalization in susceptibility.

By contrast, **resilient mice** displayed preserved (comparable to controls) volumes, FA, and efficiency in the DG, which may reflect passive maintenance, but can also indicate active adaptive processes that protect against stress-induced disruption. Such protective mechanisms may include neurogenesis, dendritic remodeling, synaptic plasticity, or glial support, which help maintain structural integrity and balanced network organization. At the network level, Resilience was associated with increased network clustering in the IAN. The BLA, IAN, and central amygdala (CeA) form the classic inhibitory gating pathway that regulates fear responses. The BLA provides excitatory input to the CeA, thereby controlling the expression of fear (34–37). In susceptible animals, elevated FA in the BLA suggests reduced microstructural complexity, which may limit the region’s flexibility under stress and contribute to stronger excitatory drive. By contrast, resilient animals displayed greater efficiency in the IAN, potentially indicating stronger inhibitory gating that prevents excessive fear responses. This pathway is further supported by our finding of lower FA in the IL and PL of resilient mice compared to controls, indicating greater microstructural flexibility—consistent with observations in stress-exposed individuals who do not develop PTSD (38). Although our data cannot directly establish increased IL/PL to IAN connectivity, the combination of reduced FA in IL/PL and increased IAN efficiency suggests a stronger influence of IL/PL over IAN within the known circuit. Given that IL and PL projections enhance inhibitory gating in the IAN during extinction, these patterns align with the extinction responsiveness observed exclusively in resilient animals (5). Together, these results indicate that resilience and susceptibility differ fundamentally in how stress shapes the amygdalar gating circuit, with resilience reflecting more effective engagement of inhibitory gating mechanisms.

By preregistering our analysis plan, we enhanced transparency, limited analytic flexibility, and strengthened the robustness of our findings. Integrating a translational behavioral model with imaging methods widely used in humans, our work bridges mechanistic insights from rodents to clinical research. New targets identified here may inform interventions such as deep brain stimulations (15) and stress inoculation training (39–40).

Our study has limitations. We cannot *fully* separate pre-existing traits from stress-induced changes, for example hippocampal differences can precede stress (30, 13). Nor can we assess reversibility, although mPFC volume recovery has been reported (41). Imaging was performed 24 hours after defeat, and some neural adaptations may emerge over longer intervals (42). Nevertheless, this timepoint aligns with the behavioral characterization window of our recently established translational model, ensuring temporal correspondence between brain and behavioral phenotyping. Including female mice in future work will be essential to examine sex-specific mechanisms. Future work should also include histological validation, as imaging differences reflect diverse processes such as neurogenesis, synaptogenesis, or myelination that cannot be distinguished without cellular analysis.

In conclusion, our study demonstrates that resilience, susceptibility, and impaired aversive learning are associated with distinct brain signatures. Resilience was characterized by preserved DG integrity, greater microstructural complexity in prefrontal subregions, unique connectivity changes (particularly in the RHR), and potentially enhanced amygdalar inhibitory gating capacity, underscoring resilience as an active rather than passive process. Meanwhile, susceptibility is consistently marked by diminished DG complexity and reduced microstructural flexibility, with a similar—though less pronounced—pattern observed in the BLA. In contrast, impaired aversive learning is associated with compromised CA1/CA2 integrity and reduced pons connectivity, underscoring the importance of hippocampal–brainstem interactions in memory consolidation (a conceptual overview is presented in Figure 6). More broadly, our results emphasize that—even in genetically identical and environmentally uniform mice—**individuals diverge in their coping strategies and neural adaptations to stress**. This individuality in brain and behavior underscores the importance of studying stress resilience not only as a clinical construct but also as a fundamental principle of how organisms cope with adversity.

**Figure 6.**
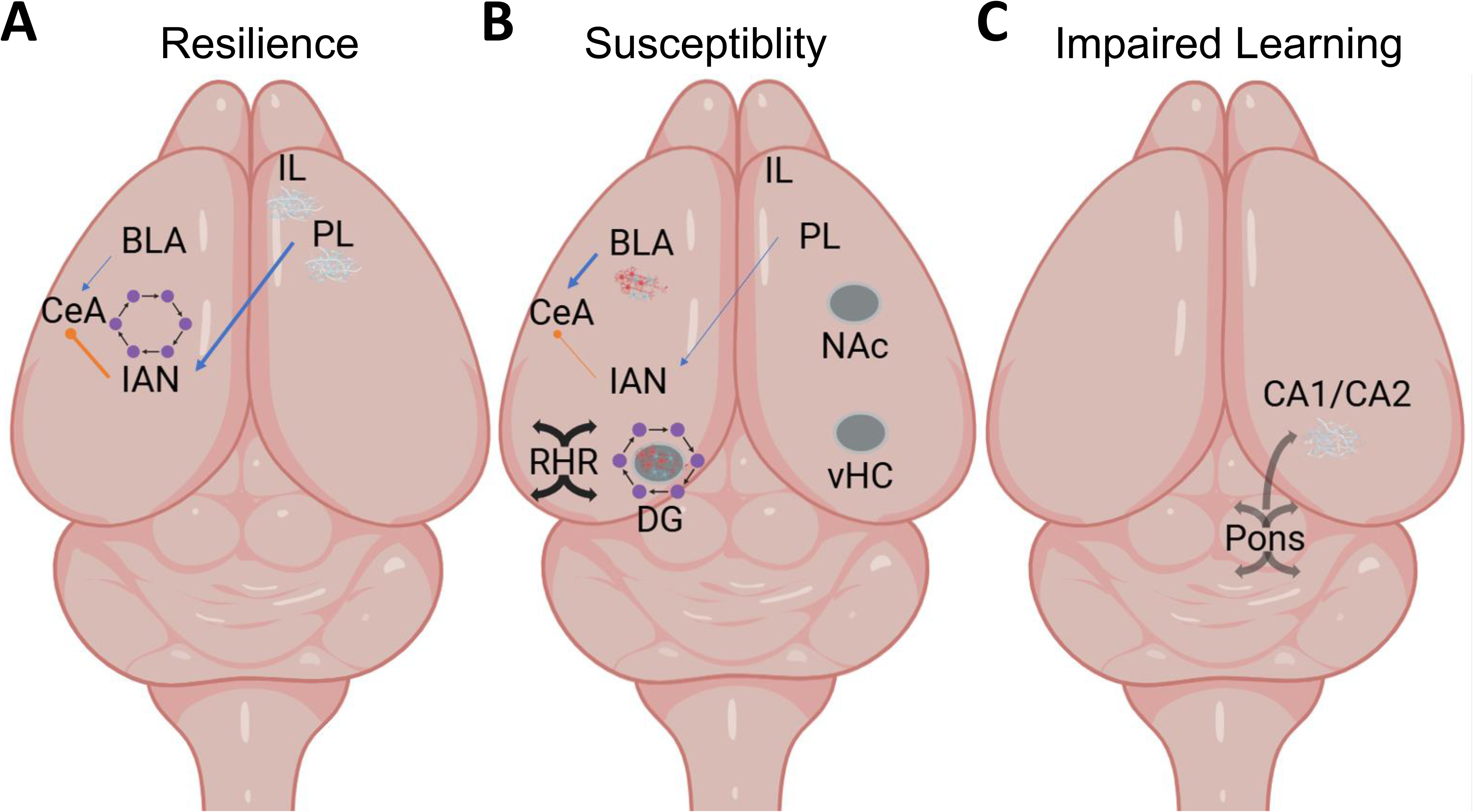
Conceptual summary of structural and network-level signatures of resilience, susceptibility, and impaired threat learning. (A) Resilience. Resilient mice show lower fractional anisotropy (FA) in infralimbic (IL) and prelimbic (PL) cortex, illustrated as more complex and branched microstructure, and higher local efficiency in the intercalated amygdala (IAN), depicted as stronger local connections. The basolateral amygdala (BLA), IAN, and central amygdala (CeA) form the inhibitory gating circuit. IL/PL provide excitatory input to the IAN, which in turn inhibits the CeA, supporting potential enhanced inhibitory gating in resilient animals. Together, these features reflect flexible prefrontal organization and stronger top-down regulation of fear. (B) Susceptibility. Susceptible mice exhibit reduced volumes in ventral hippocampus (vHC), dentate gyrus (DG), and nucleus accumbens (NAc). The DG shows the largest number of alterations, including higher FA (represented as more directionally uniform structure) and greater local efficiency, consistent with reduced microstructural complexity and less flexible network organization. The retrohippocampal region (RHR) displays stronger connectivity, and the BLA shows higher FA, suggesting diminished complexity. Within the amygdalar circuit, stronger BLA signaling toward the CeA may promote enhanced excitatory drive and elevated fear expression. (C) Impaired threat learning. Mice that fail to learn the threat-associated cue show reduced FA in CA1/CA2 of the hippocampus and weaker connectivity of the pons, illustrating disrupted hippocampal–brainstem communication. These combined alterations highlight compromised memory consolidation processes, which are essential for forming conditioned threat associations. Anatomical locations in this schematic are approximate proxies rather than precise coordinates, and the hemisphere orientation is arbitrary, as the figure is intended for conceptual illustration only.

## METHODS

### Animals

C57BL/6J male mice (n=55) weighing 22-28g at the age of 7 weeks were obtained from Janvier (France), housed individually in a temperature- and humidity-controlled facility on a 12hr light-dark cycle (23°C, 38%, lights on 8:00) with ad libitum. Procedures were performed in accordance with the European Communities Council Directive and approved by local authorities (Landesuntersuchungsamt Rheinland-Pfalz).

### Chronic social defeat

Experimental mice (Defeated n=43) were introduced into the home cage of a larger, older, and retired male breeder from the aggressor’s strain (CD-1, different individual mouse every day, pre-existing in the facility) for 10 days. After a total of 30s of attacks, animals were separated by a mesh wall overnight. During the same period, age-matched mice randomized to non-defeated controls (n=12) were handled by being placed in an empty cage for 90s before returning to home cage divided in half by a mesh wall. On the last day, all mice were housed individually in new cages to rest overnight. STST followed 24h later and was conducted between 8:30-13:30 under light conditions of 37lx.

### Social threat-safety test

The test was performed similar to Ayash and colleagues (5–6). In brief, experimental mice were introduced into a three-chambered arena, where each of the two peripheral chambers contained an empty mesh enclosure. After 6 mins of habituation phase, a 6 mins testing phase immediately followed with two larger, older, and unknown mice from different strains (CD-1 threat strain and 129/Sv unknown safe strain; pre-existing in the facility) simultaneously presented in each enclosure (Figure 1A). Using the SI index (Figure 1B) we identified three phenotypic subgroups within the Defeated group. The SI index was calculated as follows: time spent interacting with each strain during the testing phase / average time spent exploring the two empty mesh enclosures during the habituation phase.

### Tracking

Tracking of the STST was done using Ethovision software 11.0 by Noldus® (Wageningen, Netherlands). Exploration (during habituation phase of the test) and interaction (during the testing phase of the test) were scored when the nose tip of the experimental mouse was within 2cm of the area surrounding the mesh enclosures. Additionally, a blinded observer corrected for nose-tail switch errors by the software.

### Brain tissue preparation

Animals were transcardially perfused with 4% paraformaldehyde, the whole head including the skull was cut off and stored in the fixative for another 1 week at 4°C in 50mL tubes. Fixed brains were then rehydrated with 1x phosphate buffered saline (PBS). PBS was exchanged 3 times over another week to remove remaining fixative. Heads were then transferred to 15ml tubes filled with perfluorocarbon oil (Fomblin Y, Sigma Aldrich, Karlsruhe, Germany). Fomblin allows ex-vivo MRI without artefacts at air-tissue interfaces, without dehydration of the sample, and without signal from outside the head.

### Ex vivo MRI

Anatomical T2-weighted (T2w) and dMRI were performed at 7 T (BioSpec 70/20 USR, Bruker, Ettlingen, Germany) with a Tx/Rx 1H-cryoprobe and ParaVision 6.0.1 software. T2w images were acquired with a 2D-RARE sequence (repetition time TR / echo time TE=8000 ms / 33 ms, RARE factor 8, 3 averages, 64 contiguous axial slices with a slice thickness of 0.25 mm, field of view FOV=16×16 mm^2^, matrix of 200×200, band width BW=34.7 kHz, and total acquisition time TA=10:00min). Diffusion MR images were acquired with a multishot 2D spin echo EPI sequence with matching geometry and resolution (16 segments, TR=4000 ms, TE=31 ms, double sampling on, BW=100 kHz, diffusion gradient duration/separation=4.5 ms / 17 ms) with multishell diffusion encoding using 4 separate scans with one additional b=0 image and a reference frequency adjustment each (6 directions with b=100 s/mm^2^, 20 directions with b=1600 s/mm^2^, 40 directions with b=3400 s/mm^2^, 61 directions with b=6000 s/mm^2^, TA∼2:20h). High b values accounted for lower diffusivity in fixed tissue. Diffusion directions were calculated with the online tool available at http://www.emmanuelcaruyer.com/q-space-sampling.php (43), the number of points were varied linearly with diffusion wave vector q (b∼q2) i.e. with a square root dependency N(b)∼b^1/2^. T2w images were segmented into tissue probability maps of gray/white matter and cerebrospinal fluid and custom brain atlases (the atlas is available online at the study registration on Open Science Framework, see supplementary) derived from the Allen mouse brain atlas (CCFv3) were registered to the T2w and dMR images using ANTx2 (https://github.com/ChariteExpMri/antx2). dMRI was processed in mrtrix (https://www.mrtrix.org) and custom tools based on network analysis as described previously. The processing pipeline consisted of

1. Conversion of Bruker raw data into NIFTI image format
2. Denoising, Gibbs ringing removal, bias field correction, eddy current correction, and motion correction
3. Calculation of conventional diffusion tensor imaging maps of FA
4. Diffusion orientation function reconstruction using constrained spherical deconvolution
5. Connectome reconstruction using streamline tractography and SIFT2 optimization
6. Reconstruction of connectivity matrices counting the number of streamlines from atlas region to region

### Statistics

We performed group statistical comparisons (t-test, two-tailed) on

- Volume of each ROIs on T2w images
- Mean FA in each ROI
- Each connection in the connectivity matrix (whole atlas level)
- Network metrics defined for each node (=atlas region), namely local clustering coefficient and local efficiency

Comprehensive statistical information is available in figures’ legends, and supplementary tables.

## REGISTRATION AND DATA AVAILABILITY

This study has been pre-registered before brain imaging data analysis in Open Science Framework (https://doi.org/10.17605/OSF.IO/MDTH4). See supplementary for details. Raw data and secondary data are available under G-Node Infrastructure for Neuroscience (DOI 10.12751/g-node.yi7b27). The reconstructed dMRI data is too large for common open data repositories and can be shared upon request.

## CODE AVAILABILITY

Code is available at https://github.com/ChariteExpMri/dmri_pipe.

## ACKNOWLEDGEMENTS

Funding to S.A. was provided by the Einstein Foundation (Einstein Independent Researcher Grant). Funding to M.B.M was supported by the German Research Foundation (DFG, CRC 1193, Z02), the Boehringer Ingelheim Foundation (Individual phenotyping and high-resolution automated behavioral analysis), and by Leibniz Competition Funds (Leibniz ScienceCampus NanoBrain, W71/2022). Funding to SM, SPK, and PBS was provided by the German Federal Ministry of Education and Research (BMBF) under the ERA-NET NEURON scheme (01EW2305), and the German Research Foundation (DFG, project BO 4484/2-1, Project-ID 424778381-TRR 295 ReTune and EXC-2049-390688087 NeuroCure). Imaging experiments were supported by Charité 3R – Replace | Reduce | Refine. Figure 1A and Figure 6 schematic depictions are adapted from icons by BioRender.com. The funding sources were not involved in the study. We thank Ulf Toelch for a valuable discussion on data analysis.

## AUTHOR CONTRIBUTION

S.A., M.B.M., and U.S. conceived the idea; S.A. and P.B.S registered the study; S.A. planned and carried out the experiments and performed the behavioral analysis; M.F. and E.C. provided technical support with the behavioral experiments; S.M. and P.B.S. supported S.A. with acquiring the brain imaging data; J.A. pre-processed the brain imaging data; S.P.K. and P.B.S. analyzed the brain imaging data; S.A., P.B.S., and S.P.K. interpreted the results; U.S., M.B.M., and OT supported with interpretations of the results; S.P.K. created the figures; S.A. wrote the manuscript; M.B.M, P.B.S., and U.S. supported with writing the manuscript. All authors reviewed and approved the final submitted manuscript.

