## Supplementary Material for "Network Adaptability Governs Resilience and Susceptibility to Social Defeat"

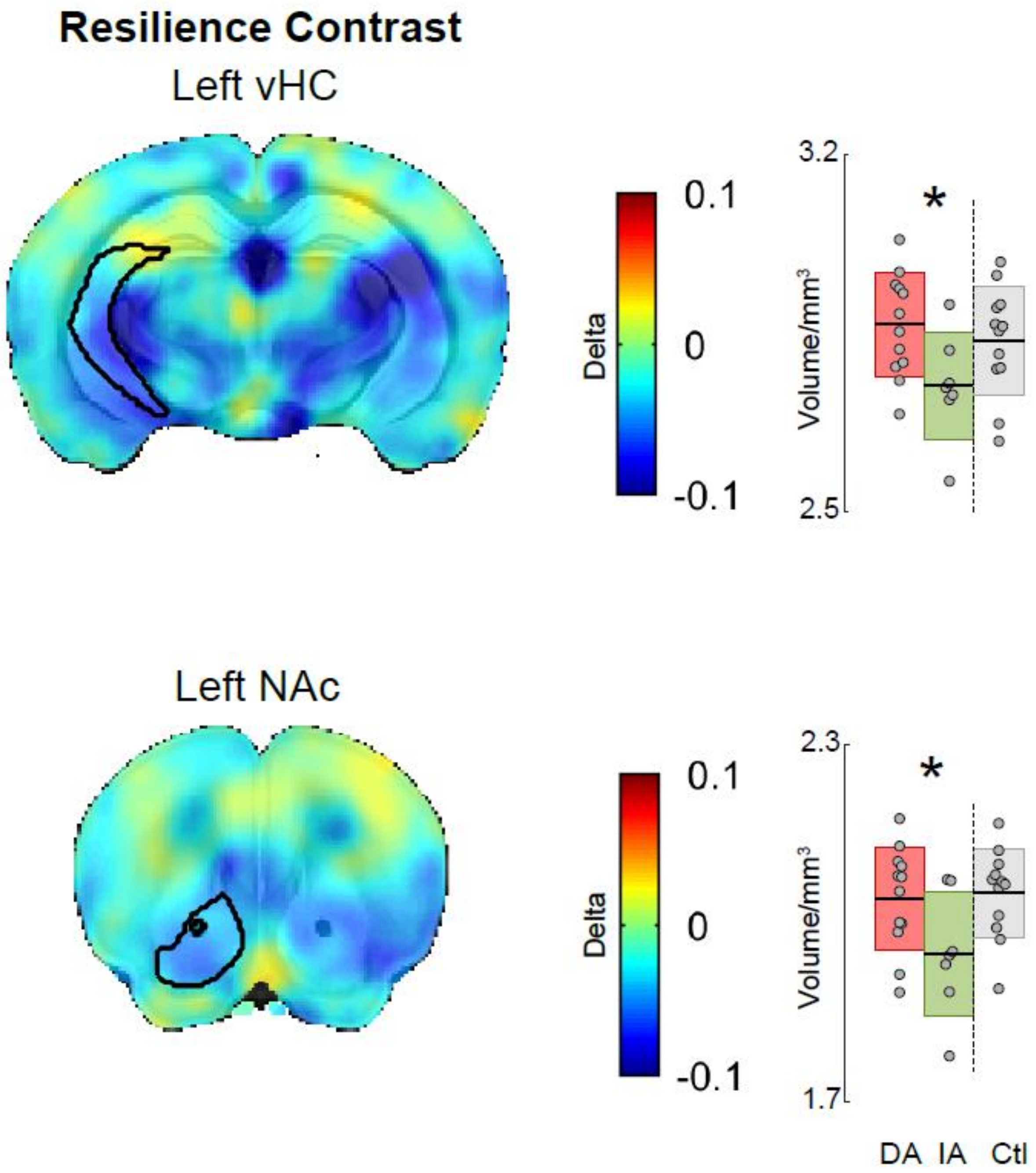


**Figure 1. Region wise volume in resilience-associated differences.** Ventral hippocampus (vHC) and nucleus accumebns (NAc) are significantly larger in resilient discriminating-avoider (DA; n=12) compared to susceptible indiscriminate-avoider (IA; n=7). Data from control animals is displayed for visualization purposes only and was excluded from statistical analysis. Brain maps depict the differences quantified in the respective box plot, the region of interest is marked in black, color bars indicate relative volume differences, with volumes normalized to the Allen Brain Atlas and compared between groups. Box plots show mean±SD., t-test, two-tailed, vHC (p-value=0.02,

Mean of DA=2.92 and SD=0.11, Mean of IA=2.78 and SD=0.12, t-value=2.47, df=17), and NAc (p-value=0.05, Mean of DA=2.04 and SD=0.09, Mean of IA=1.95 and SD=0.10, t-value=2.11, df=17).

**Table 1A: Region-wise volume. Indiscriminate-avoiders versus controls**

| Brain Region | P-value | T-value | Mean 1 (SD1) | Mean 2 (SD2) |
|---|---|---|---|---|
| Nucleus accumbens (left) | 0.02# | -2.52 | 1.95 (0.10) | 2.05 (0.07) |
| Nucleus accumbens (right) | 0.08 | -1.83 | 1.94 (0.10) | 2.02 (0.09) |
| Dentate gyrus (right) | 0.04# | -2.24 | 2.92 (0.14) | 3.09 (0.16) |
| Dentate gyrus (left) | 0.31 | -1.05 | 3.02 (0.13) | 3.09 (0.16) |

**Table 1B: Region-wise volume. Discriminating-avoiders versus controls**

| Brain Region | P-value | T-value | Mean 1 (SD1) | Mean 2 (SD2) |
|---|---|---|---|---|
| Nucleus accumbens (left) | 0.70 | -0.27 | 2.04 (0.09) | 2.05 (0.07) |
| Nucleus accumbens (right) | 0.41 | 0.84 | 2.05 (0.09) | 2.02 (0.09) |
| Dentate gyrus (right) | 0.49 | -0.70 | 3.05 (0.10) | 3.09 (0.16) |
| Dentate gyrus (left) | 0.72 | -0.36 | 3.07 (0.09) | 3.09 (0.16) |

**Table 1. Regions-wise volume. (A) Indiscriminate-avoider versus controls.** Nucleus accumbens and dentate gyrus are significantly (#) smaller in volume in susceptible indiscriminate-avoiders (n=7) compared to non-defeated controls (n=12). **(B) Discriminating-avoiders versus controls.** No significant differences observed in volume between resilient discriminating-avoiders (n=12) and non-defeated controls (n=12). T-test, two-tailed, mean 1 and SD 1 belong to the first group in the title i.e. (A) indiscriminate-avoiders, (B) discriminating-avoiders. (A) df=17, (B) df=22.

**Table 2A: Diffusion magnetic resonance imaging fractional anisotropy. Indiscriminate-avoiders versus Controls**

| Brain Region | P-value | T-value | Mean 1 (SD1) | Mean 2 (SD2) |
|---|---|---|---|---|
| Dentate gyrus (left) | 0.03# | 2.39 | 0.22 (0.02) | 0.20 (0.01) |
| Dentate gyrus (right) | 0.25 | 1.19 | 0.22 (0.02) | 0.21 (0.01) |
| Basolateral amygdala (right) | 0.01# | 2.83 | 0.23 (0.04) | 0.19 (0.01) |

| Brain Region | P-value | T-value | Mean 1 (SD1) | Mean 2 (SD2) |
|---|---|---|---|---|
| Basolateral amygdala (left) | 0.52 | 0.65 | 0.21 (0.03) | 0.21 (0.02) |
| Prelimbic cortex (left) | 0.16 | -1.46 | 0.21 (0.02) | 0.23 (0.03) |
| Prelimbic cortex (right) | 0.16 | 1.45 | 0.24 (0.02) | 0.22 (0.02) |
| Infralimbic cortex (right) | 0.25 | 1.19 | 0.20 (0.02) | 0.19 (0.03) |
| Infralimbic cortex (left) | 0.41 | -0.84 | 0.19 (0.02) | 0.20 (0.03) |

**Table 2B: Diffusion magnetic resonance imaging fractional anisotropy. Discriminating-avoiders versus Controls**

| Brain Region | P-value | T-value | Mean 1 (SD1) | Mean 2 (SD2) |
|---|---|---|---|---|
| Prelimbic cortex (right) | 0.01# | -2.82 | 0.19 (0.02) | 0.22 (0.02) |
| Prelimbic cortex (left) | 0.81 | -0.24 | 0.23 (0.02) | 0.23 (0.03) |
| Infralimbic cortex (right) | 0.01# | -2.78 | 0.16 (0.02) | 0.19 (0.03) |
| Infralimbic cortex (left) | 0.57 | -0.58 | 0.20 (0.03) | 0.20 (0.03) |
| Dentate gyrus (right) | 0.10 | -1.74 | 0.20 (0.01) | 0.21 (0.01) |
| Dentate gyrus (left) | 1.00 | 0.00 | 0.20 (0.01) | 0.20 (0.01) |
| Basolateral amygdala (left) | 0.49 | 0.71 | 0.21 (0.02) | 0.21 (0.02) |
| Basolateral amygdala (right) | 0.81 | -0.25 | 0.19 (0.01) | 0.19 (0.01) |

**Table 2. Diffusion magnetic resonance imaging fractional anisotropy. (A) Indiscriminate-avoider versus controls.** Dentate gyrus and basolateral amygdala are significantly (#) larger in fractional anisotropy in susceptible indiscriminate-avoiders (n=7) compared to non-defeated controls (n=12). **(B) Discriminating-avoiders versus controls.** Prelimbic- and infralimbic cortex are significantly (#) smaller in resilient discriminating-avoiders (n=12) compared to non-defeated controls (n=12). T-test, two-tailed, mean 1 and SD 1 belong to the first group in the title i.e. (A) indiscriminate-avoiders, (B) discriminating-avoiders. (A) df=17, (B) df=22.

**Table 3. Diffusion magnetic resonance imaging connectivity. (A) Chronic stress exposure-associated differences.** A total of 24 differential connectivities involving 29 brain regions (irrespective of hemispheres) are identified between defeated animals (n=43) and handled controls (n=12). Out of the 24, 13 are less connected in defeated. Pallidum (dorsal, ventral, and medial region) has the largest number of differential connectivities all of which are less connected in defeated. **(B) Threat learning-associated differences.** A total of 37 differential connectives involving 39 brain regions (irrespective of hemispheres) are identified between animals that don't conditionally learn to avoid the threatening strain (non-avoider; n=24) and avoiders (n=19) of threat. Out of 37, 22 are less connected in non-avoiders. Pons has the largest number of differential connectivities all of which are less connected in non-avoiders. **(C) Resilience-associated differences.** A total of 56 differential connectivities involving 47 brain regions (irrespective of hemispheres) are identified between resilient discriminating-avoiders (n=12) and susceptible indiscriminate-avoiders (n=7). Out of the 56, 28 are less connected in discriminating-avoiders. Retrohippocampal region has the largest number of differential connectivities all of which are less connected in discriminating-avoiders except for the connection with ventral tegmental area and the basomedial amygdalar area. Group differences in connectivity strength, t-test, two-tailed, mean 1 and SD 1 belong to the first group in the title i.e. (A) defeated, (B) non-avoiders, and (C) discriminating-avoiders. (A) df=53, (B) df=41 , (C) df=17.

**Table 4A: Graph theory: clustering coefficient. Discriminating-avoiders versus Controls**

| Brain Region | P-value | T-value | Mean 1 (SD1) | Mean 2 (SD2) |
|---|---|---|---|---|
| Dentate gyrus (left) | 0.62 | -0.50 | 5.69 (0.43) | 5.77 (0.37) |
| Dentate gyrus (right) | 0.58 | -0.56 | 5.95 (0.45) | 6.04 (0.33) |
| Intercalated amygdala nucleus (left) | 0.04# | 2.14 | 6.12 (0.75) | 5.43 (0.84) |
| Intercalated amygdala nucleus (right) | 0.50 | 0.69 | 5.94 (0.69) | 5.69 (1.05) |

**Table 4B: Graph theory: efficiency. Discriminating-avoiders versus Controls**

| Brain Region | P-value | T-value | Mean 1 (SD1) | Mean 2 (SD2) |
|---|---|---|---|---|
| Dentate gyrus (left) | 0.67 | -0.44 | 6.62 (0.43) | 6.69 (0.39) |
| Dentate gyrus (right) | 0.52 | -0.65 | 6.78 (0.46) | 6.88 (0.33) |
| Intercalated amygdala nucleus (left) | 0.03# | 2.34 | 6.24 (0.71) | 5.52 (0.79) |
| Intercalated amygdala nucleus (right) | 0.45 | 0.77 | 6.03 (0.70) | 5.75 (1.07) |

**Table 4C: Graph theory: clustering coefficient. Indiscriminate-avoiders versus Controls**

| Brain Region | P-value | T-value | Mean 1 (SD1) | Mean 2 (SD2) |
|---|---|---|---|---|
| Dentate gyrus (left) | 0.06 | 1.99 | 6.55 (0.28) | 6.26 (0.31) |
| Dentate gyrus (right) | 0.67 | -0.44 | 6.19 (0.30) | 6.25 (0.25) |
| Intercalated amygdala nucleus (left) | 0.34 | -0.99 | 6.13 (1.51) | 6.68 (0.94) |
| Intercalated amygdala nucleus (right) | 0.75 | 0.33 | 6.68 (0.30) | 6.56 (1.01) |

**Table 4D: Graph theory: efficiency. Indiscriminate-avoiders versus Controls**

| Brain Region | P-value | T-value | Mean 1 (SD1) | Mean 2 (SD2) |
|---|---|---|---|---|
| Dentate gyrus (left) | 0.04# | 2.17 | 7.67 (0.29) | 7.31 (0.37) |
| Dentate gyrus (right) | 0.82 | -0.23 | 7.28 (0.34) | 7.32 (0.29) |
| Intercalated amygdala nucleus (left) | 0.30 | -1.06 | 6.14 (1.51) | 6.72 (0.10) |
| Intercalated amygdala nucleus (right) | 0.73 | 0.35 | 6.76 (0.29) | 6.62 (1.02) |

**Table 4. Graph theory. (A) Clustering coefficient: discriminating-avoiders versus control.** Intercalated amygdala nucleus displays significantly (#) greater clustering coefficient in resilient discriminating-avoiders (n=12) compared to non-defeated controls (n=12). **(B) Efficiency: discriminating-avoiders versus controls.** Intercalated amygdala nucleus displays significantly (#) greater efficiency in resilient discriminating-avoiders compared to non-defeated controls. **(C) Clustering coefficient: indiscriminate-avoiders versus controls.** Dentate gyrus displays a trend in greater clustering coefficient in susceptible indiscriminate-avoiders (n=7) compared to controls (n=12). **(D) Efficiency: indiscriminate-avoiders versus controls.** Dentate gyrus displays significantly (#) greater efficiency in susceptible indiscriminate-avoiders compared to non-defeated controls. T-test, two-tailed, mean 1 and SD 1 belong to the first group in the title i.e. (A-B) discriminating-avoiders, (C-D) indiscriminate-avoiders, (A-B) df=22, (C-D) df=17.

REGISTRATION

This study has been pre-registered before data analysis in Open Science Framework (https://doi.org/10.17605/OSF.IO/MDTH4). We reduced the Allen Mouse Brain atlas to 44 regions out of 372. The atlas is deposited in the pre-registration. Differences between our report here and our pre-registration are: 1. In addition to the 10 regions of interest (ROIs) we report here, we performed analysis on a smaller set of ROIs (three regions), namely basolateral amygdala (BLA), hippocampus (HC), and ventral tegmental area (VTA) that we don’t report in our paper. Our decision is motivated by our previous transcriptome analysis on the same mouse model (Ayash et al., 2023a) where we targeted key regions of the fear circuitry, namely medial prefrontal cortex (mPFC), ventral HC (vHC), and basolateral amygdala (BLA). In the set of the 10 ROIs, all subregions of those three regions are included in addition to three more, namely VTA, nucleus accumbens (NAc), and hypothalamic medial zone (HMZ) that literature strongly suggests are implicated in response to chronic social defeat (CSD). Thus, we think the 10 ROIs are more informative than the three ROIs and build on the findings from the previous transcriptome analysis. 2. In the pre-registration, we define stress effect as control animals compared to susceptible indiscriminate-avoiders and resilience discriminating-avoiders i.e. we don't mention non-avoiders of threat. Stress effect should include all animals that underwent CSD independent if they learn to socially avoid the aggressors’ strain or not. Accordingly, in our paper all defeated animals are included in the comparison to the control group to assess stress effect. 3. Multiple analyses we stated in the pre-registration that we will perform that we don’t report here. This includes: whole brain volume, cluster peak for voxel-based, mean and radial diffusivity, modularity, transitivity, and assortivity. To keep the report focused on the most relevant findings, we decided not to report the results of these analyses. 4. In the pre-registration, we state that we will perform spearman correlation between MRI findings and social avoidance behavior. We didn’t perform this analysis as it would be repetitive to the work Anacker et al., 2016 did (see reference section in the paper). We decided to focus on analysis that has not been done before. Moreover, our mouse model conceptualizes social avoidance in a different way to the model employed by all other literature employing similar experiments, thus it doesn’t contribute to our study to perform this analysis. 5. Finally, in the pre-registration we state we will perform the volume analysis as exploratory on whole atlas level, we later decided to perform it as a hypothesis driven analysis on the 10 ROIs and restrict exploratory analysis to connectivity assessment and graph theory measurements.
